# Quantifying changes in pain sensitivity using reproducible transcutaneous optogenetic stimulation in behaving mice

**DOI:** 10.1101/2024.12.13.628414

**Authors:** Yu-Feng Xie, Christopher Dedek, Steven A. Prescott

## Abstract

Optogenetics provides an unprecedented opportunity to delineate how different somatosensory afferents contribute to sensation, including pain. Afferents expressing channelrhodopsin-2 (ChR2) can be selectively activated by transcutaneous photostimuli applied to behaving mice. Despite care taken to precisely target expression of ChR2 to specific cell types, imprecise photostimulation has hindered quantitative optogenetic-based behavioral testing. Here, we used a robot to reproducibly apply transcutaneous optogenetic stimuli to the hind paw of mice while measuring nocifensive withdrawal. Different photostimulus waveforms (pulses and ramps) and response metrics (threshold and latency) were compared in mice expressing ChR2 in all afferents (Advillin-ChR2) or preferentially in nociceptors (Na_V_1.8-ChR2). CFA-induced inflammation caused withdrawal from ramped photostimuli to become faster relative to baseline and to vehicle-injected controls whereas pulsed photostimuli revealed a modest increase in threshold. Analgesia caused by Na_V_1.7 and 1.8 channel blockade was evident in both testing protocols. Overall, ramp-based testing was more effective and more efficient (i.e. required less time and total stimulation) than pulse-based testing. Electrophysiological measurements confirmed that inflammation increases nociceptor excitability without affecting phototransduction, and suggests that withdrawal latency depends on the number of nociceptors activated rather than how strongly each nociceptor is activated. Consistent with changes described in nociceptor somata, the behavioral consequences of peripherally blocking different voltage-gated sodium (Na_V_) channels showed that nociceptor axons normally rely on Na_V_1.8 but upregulate Na_V_1.7 after inflammation, with important clinical implications for drug efficacy. Collectively, these results demonstrate the utility of optogenetic pain testing when reproducibly delivered and strategically designed photostimuli are used.

## INTRODUCTION

Optogenetics has revolutionized neuroscience by enabling specific neurons to be activated or inhibited using light [17]. Specificity is achieved by targeting expression of light-sensitive actuators like ChR2 to genetically defined cell types [26; 30; 35; 47]. Photostimuli are typically applied as pulses intended to evoke one spike per pulse [33]. This allows photostimulus pulse rate to control neuron firing rate, but it comes at the expense of evoking unnaturally synchronized spiking across activated neurons. Synchrony can be mitigated by adjusting photostimulus kinetics [56] but use of non-pulsed photostimuli is rare [2; 28; 56]. Furthermore, even if each photostimulus pulse evokes one spike per neuron, the intensity and spatial distribution of applied light affects the number of activated neurons. Controlled, reproducible photostimulation is thus necessary to explore population-level coding. Indeed, precise photostimulation using optrodes [20] or multiphoton excitation [36] has enabled phototagging [11] and efficient circuit mapping [24] where the extent of neuronal activation must be controlled. New optogenetic applications promise to emerge from improvements in photostimulus design and delivery, which may especially benefit investigation of somatosensory processing.

Optogenetics has already been applied along the entire somatosensory neuraxis [14; 53], including at early stages of sensory transduction [29]. Cutaneous afferents can be photostimulated non-invasively through the skin but spatiotemporal targeting of transcutaneous photostimuli has been crude. Past studies measuring withdrawal from transcutaneous photostimuli have relied exclusively on pulses [1; 5-7; 9; 12; 15; 18; 23; 31; 32; 44; 45; 48; 50], mostly applied by handheld fiber optic cables, with some exceptions [16; 44]. Uncontrolled variability in stimulation and imprecise response measurement severely limit quantitative pain testing. This motivated us to develop a robot, RAMalgo, to reproducibly deliver optogenetic (and other) stimuli and precisely measure paw withdrawal [16]. Notably, poor reproducibility is not limited to optogenetic testing; applying any stimulus by hand and scoring responses by eye leads to low inter-tester reliability [57]. Minimizing this technical variability is crucial for studying meaningful biological variability and for resolving subtle changes in pain sensitivity.

Here, we used RAMalgo to compare responses evoked by different photostimulus waveforms. Pulsed photostimuli activate neurons synchronously whereas ramped photostimuli are more likely to cause asynchronous activation, and may not even activate neurons whose spike initiation requires rapid depolarization [56]. This likely translates to different photostimulus waveforms activating different afferents with different patterns, where some input patterns evoke spinally-mediated reflexive withdrawal whereas others allow time for supraspinal circuits to contribute. It is, therefore, notable that changes in pain sensitivity were more readily evident from ramp-based testing. Electrophysiological data suggest that changes in withdrawal from ramped photostimuli reflect the number of activated afferents. By selectively blocking Na_V_1.8 or Na_V_1.7 channels in nociceptive axon terminals of naïve and inflamed mice, we also show that inflammation shifts the relative contribution of these Na_V_ isoforms to axon excitability. With the right tools and protocols, optogenetic testing can evidently detect changes in pain sensitivity while also helping to delineate the contribution of different afferent types. These results especially highlight importance of the photostimulus waveform.

## METHODS

### Animals

All procedures were approved by the Animal Care Committee at The Hospital for Sick Children (protocol #65769) and conducted in compliance with guidelines from the Canadian Council on Animal Care. Homozygous Ai32 mice (Jax #012569), which express ChR2(H134R)-eYFP in a Cre-dependent manner, were crossed with advillin-Cre mice provided by Fan Wang (equivalent to Jax #032536) to generate mice expressing ChR2 in all afferents [55] or with Na_V_1.8-Cre mice to create mice expressing ChR2 preferentially in nociceptive afferents [3]. Two Na_V_1.8-Cre lines were used, namely Tg(Scn10a-cre)1Rkun mice provided by Rohini Kuner and B6.129(Cg)-*Scn10a^tm2(cre)Jwo^*/TjpJ (Jax # 036564) mice created by John Wood. The resulting mice are referred to as Na_V_1.8-ChR2 but are designated RK or JW. Expression of ChR2 is not strictly limited to nociceptors in Na_V_1.8-Cre mice since Na_V_1.8 is also expressed in C-LTMRs and in some A fibers [46] (see Discussion).

### Behavioral testing

Behavioral testing was performed on mice of either sex at 8-12 weeks of age. No sex differences were observed (see Results) and data were therefore pooled. Mice were acclimated to the testing environment for 60 min/day for two days before the testing, and for 60 min prior to each test session. Behavioral tests were conducted with a Reproducible Automated Multimodal algometer, RAMalgo (Robotic Algometry Inc., Toronto) [16]. Mice were viewed from below by video. Aiming was controlled by joystick so that the human tester remained distant from the mice throughout testing. Once a trial was initiated by the tester, all subsequent steps (i.e. stimulation and response measurement) were automated. The red light, whose reflectance level is used to measure paw withdrawal (see Fig. 1), was set at 0.3 V with a termination threshold of 0.9.

**FIGURE 1.**
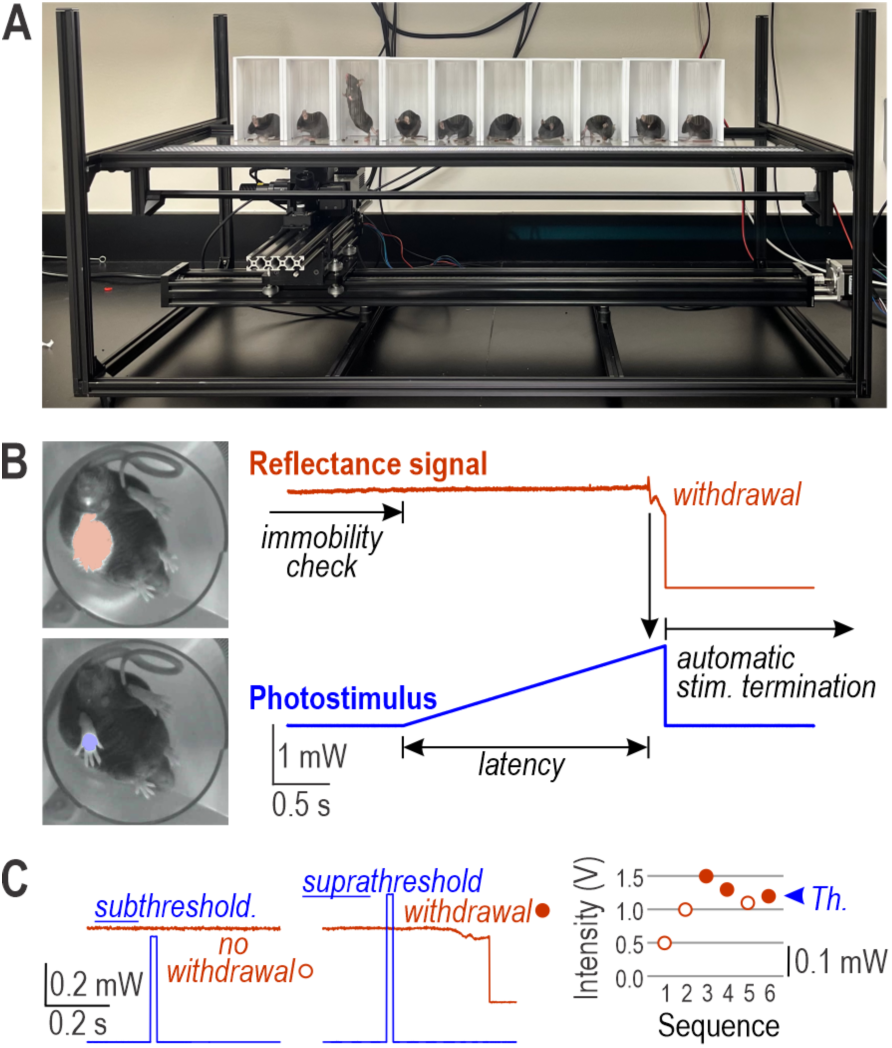
Quantitative transcutaneous optogenetic stimulation with different photostimulus waveforms. **(A)** RAMalgo with 10 mice on the testing platform. The stimulator is located below the platform, mounted on motorized actuators for joystick-controlled aiming. Components for mechanical stimulation are removed to give a clearer view of photostimulation components. Mice undergo one trial each in rapid succession before starting a second round of trials, thus ensuring each mouse has a long inter-stimulus interval while minimizing the total time taken to test all mice. **(B)** Ramped photostimulation (see also Video 1). The target paw is first illuminated with red light whose reflectance off the paw is measured by a photodetector sampling at 1 kHz (red trace). Photostimulation with blue light (bottom) is initiated automatically if the paw is deemed immobile (based on stability of the red reflectance signal) during the user-defined pre-stimulus period. Paw withdrawal triggers abrupt changes in the reflectance signal, enabling precise measurement of withdrawal latency and automatic termination of stimulation. Photos show frames captured from video. Blue photostimulation is restricted to a spot centered on the paw within the larger area of red illumination; for illustrative purposes, each color is shown separately with saturated pixels pseudocolored in blue or red. **(C)** Pulsed photostimulation. Red light is used like in B, but blue photostimulation is applied as a 20 ms-long pulse. Sample trials are shown for a pulse at subthreshold intensity (left), which evokes no withdrawal (open circle), and at suprathreshold intensity (right), which evokes a withdrawal (filled circles). Inset shows how pulse intensity is incremented in 0.5 V steps until withdrawal occurs, then decremented in 0.2 V steps until withdrawal does not occur, and then incremented in 0.1 V steps until withdrawal occurs. The final intensity is considered threshold (Th.).

Blue light for optogenetic stimulation was focused to a spot 3.5 mm in diameter on the left hind paw. A 1 V control signal yields 0.225 mW/mm^2^ of blue light on the target as measured with a s170C photodiode attached to a PM100D optical power meter (Thorlabs) whose output was normalized by the spot’s area. Photostimuli were applied as a pulse or ramp. For pulsed photostimuli, light was applied for 20 ms at the user-defined intensity. A single pulse was applied per trial. An up-down titration method was used to determine threshold: pulse intensity was increased in 0.5 V steps until paw withdrawal occurred, then decreased in 0.2 V steps until withdrawal did not occur, and finally increased in 0.1 V steps until withdrawal occurred again, with the last stimulus intensity designated as threshold. For each mouse, the average threshold from three titration sequences is reported per test session. For ramped photostimuli, light intensity was ramped to 4 V over 15 s; stimulation stops automatically once withdrawal is detected. The average withdrawal latency from three ramps is reported for each test session. Data from each trial were saved to electronic files along with metadata and video. A test session was conducted at baseline and again 24 and 48 hours after injecting CFA or saline. When drug (PF-24 and/or PF-71) or its vehicle was injected, another test session was conducted starting 30 min after injection.

#### CFA injection

To induce inflammation, 20 µl of complete Freund’s adjuvant (CFA, Sigma, F5881) was injected subdermally into the left hind paw shortly after baseline testing. Mice were lightly anesthetized with isoflurane (3.5%) during injection. Control mice received 20 µl of vehicle, namely phosphate-buffered saline (PBS). Mice were randomly allocated to CFA or saline groups but the experimenter was not blinded during testing since robotic testing precludes biased stimulation or response measurement.

#### Drug administration

PF-05089771 (PF-71, 300 nM, Alomone) and PF-01247324 (PF-24, 10 µM, Sigma) were first dissolved in DMSO and then diluted to final concentration with PBS. 20 µl of either drug, both drugs together, or vehicle (equivalent DMSO diluted in PBS) was injected subdermally into the left hind paw under light anesthesia with isoflurane. Mice were randomly allocated to drug or vehicle groups.

### Acute isolation of dorsal root ganglia (DRG) neurons

DRG neurons were obtained from adult (8-12 week-old) Na_V_1.8-ChR2 mice as previously described [54]. Mice were anesthetized with isoflurane (3.5%) and perfused intracardially with ice-cold HBBS (Ca/Mg free, LifeTech 14170112) supplemented with (in mM) 15 HEPES, 28 Glucose, and 111 sucrose; pH was adjusted with NaOH to 7.3-7.4; osmolarity was 312-319 mOsm. Lumbar DRGs (left L2-5 for naïve mice and L4 only for CFA-injected mice) were collected. After digestion with papain (Worthington Biochemical) and collagenase (Worthington Biochemical)/dispase II (Sigma), DRGs were mechanically triturated, plated onto poly-D lysine-coated coverslips, and incubated in Neurobasal™ media (Gibco 21103-049) supplemented with 1% fetal bovine serum (FBS), B-27™ supplement (Thermo Fisher 17504-044) and 0.5 mM L-Glutamine (Gibco 25030-081) for 2 hours. After this, fresh media was added to the wells after removal of unattached cells and cells were maintained in a 5% CO_2_ incubator at 37°C until recording.

### Electrophysiological recording

DRG neurons were tested 2-8 h after plating to avoid changes in Na_V_ expression [54] and ChR2 expression [56] reported to occur after culturing. Coverslips were transferred to a recording chamber perfused with oxygenated (5% CO_2_:95% O_2_) artificial cerebrospinal fluid containing (in mM) 126 NaCl, 2 KCl, 2 CaCl_2_, 1.25 NaH_2_PO_4_, 26 NaHCO_3_, 2 MgCl_2,_ and 10 glucose at room temperature. Neurons were visualized with a Zeiss AxioExaminer microscope with a 40x, 0.75 NA water immersion objective (N-Achroplan, Zeiss) and IR-1000 Infrared CCD camera (Dage-MTI). A blue LED (Thorlabs, M455L4) was attached to the epifluorescent port through a collimator; this is the same LED used on RAMalgo. A long-pass filter (OG590) was placed in the transmitted light path to prevent unintended ChR2 activation during recording. Small neurons (diameter <25 μm) expressing YFP were targeted for patching. Whole cell patch clamp recordings were performed using a pipette solution containing (in mM) 140 K-gluconate, 2 MgCl_2_, 10 HEPES, 0.2 EGTA, 3.8 Na-ATP, and 0.4 Na-GTP (pH ∼7.3, osmolarity ∼300 mOsm). Series resistance was compensated >70%. Signals were amplified with an Axopatch 200B amplifier (Molecular Device), low-pass filtered at 2 kHz, digitized with a Power1401 DAQ (Cambridge Electric Design, CED), and recorded at 10 kHz using Signal v7 (CED). All reported voltages are corrected for junction potential (-15 mV).

After determining resting membrane potential, voltage was adjusted to -70 mV to characterize excitability. Spikes were evoked by a series of 1-s-long depolarizing current steps. Rheobase, defined as the minimal current required to evoke a spike, was determined and current steps were then applied at 0.5x increments of rheobase. Firing >2 spikes in response to any stimulus intensity qualifies a neuron as repetitive spiking; transient spiking is defined as consistently firing ≤2 spikes. Spike threshold was defined as voltage where dV/dt exceeds 5 mV/ms. Optogenetic stimuli were applied in two ways. To measure ChR2 current, the neuron was voltage clamped at -70 mV while applying blue light as either a series of 1 s-long steps incremented at 0.2 V (max 2 V) or a 15 s-long ramp (max 4 V). The latter stimulus was also applied in current clamp to characterize optogenetically evoked spiking. Light intensity at the neuron was measured using a s170C photodiode attached to a PM100D optical power meter (Thorlabs); 1 V = 78 mW/mm^2^.

### Statistical analysis

Data were analysed with SigmaPlot 11 (SYSTAT). Normality of distributions was tested with the Shapiro-Wilk test. Data are summarized as mean ± SEM or as median, interquartile range (IQR, i.e. 25^th^-75^th^ percentile) for normally and non-normally distributed data, respectively. Box-and-whisker plots show the median (thick bar), IQR (gray box), and 5^th^-95^th^ percentile (whiskers). Appropriate parametric or non-parametric tests were applied. Tests and degrees of freedom are identified when reporting results. Effect size was quantified as Hedges’ *g*, which is Cohen’s *d* (the difference in means divided by pooled standard deviation) corrected for different sample sizes.

## RESULTS

To minimize stress due to handing [22], mice were transferred from their home cage in clear plastic tubes that are placed upright on the platform and slid into an opaque cubicle for testing (**Fig. 1A**). Mice were viewed from below by video (**Fig. 1B insets**). The stimulator is mounted on motorized actuators so that it can be aimed autonomously (using AI) or by joystick; in either case, the tester remains distant from the mice throughout testing. The target paw was then illuminated with red light while monitoring reflectance of that light off the paw; paw withdrawal causes an abrupt change in reflected light (**Fig. 1B**) from which withdrawal latency can be precisely measured [16]. Photostimulation was applied to a spot 3.5 mm in diameter. **Figure 1B** illustrates a typical response to ramped photostimulation; **Video 1** shows the full response for comparison with an unresponsive ChR2-negative mouse (**Video 2**). **Figure 1C** shows sample responses to pulsed photostimulation, illustrating the absence (left) or presence (right) of withdrawal depending on pulse intensity. Threshold is identified using an up-down titration method (inset). All data (including video) and metadata (date, time, stimulus parameters, etc.) were recorded to computer.

### Comparison of ramp- and pulse-based testing protocols

To quantify changes in pain sensitivity, we compared withdrawal from transcutaneous optogenetic stimulation applied to the hind paw at baseline and for two days after injecting complete Freund’s adjuvant (CFA) to induce inflammation. Different photostimulus waveforms (pulses and ramps) and response metrics (threshold and latency) were compared, starting first with Na_V_1.8_JW_-ChR2 mice, which express ChR2 predominantly in nociceptors.

Inflammation was expected to reduce threshold (i.e. increase sensitivity) but thresholds measured using pulsed photostimuli did not differ between inflamed and control mice one day post-treatment (*T*_11_ = 1.206, p = 0.25, unpaired t-test) and threshold was actually *higher* in inflamed mice by day 2 (*T*_11_ = 2.712, *p* = 0.020) (**Fig. 2A left**), although the CFA-induced increase relative to baseline did not reach significance (*T*_8_ = -1.892, *p* = 0.095, paired *t*-test) (**Fig. 2A right**). Saline injection caused a similar increase on day 1 (*T*_3_ = -3.576, *p* = 0.037, paired t-test). On the other hand, analgesia induced by injecting PF-24 and PF-71 into the paw (to block Na_V_1.8 and Na_V_1.7 channels at the photostimulation site) was expected to increase threshold, which it did (*T*_8_ = -3.502, *p* = 0.010, paired *t*-test). Withdrawal latency from pulsed photostimuli was not expected to change after CFA because latency depends on pulse intensity relative to threshold [16], and pulse intensity was adjusted to threshold in each condition. As expected, withdrawal latency to threshold-intensity pulses did not differ between baseline (396 ± 50 ms) and CFA (385 ± 31 ms) (*T*_4_ = 0.477, *p* = 0.66, paired *t*-test). Withdrawal from pulsed photostimulation occurs 100s of milliseconds after the 20 ms pulse is complete, which is slower than responses to strong suprathreshold stimuli, which take as little as 20 ms, but is still much faster than responses to ramped photostimuli. Withdrawal latencies measured in the same mice using ramps were significantly shorter in inflamed mice compared to controls (day 1: *T*_11_ = -3.734, *p* = 0.003; day 2: *T*_11_ = -2.937, *p* = 0.014, unpaired t-tests) (**Fig. 2B left**), and withdrawal was significantly faster after inflammation compared to baseline (*T*_8_ = 4.280, *p* = 0.003, paired *t*-test) (**Fig. 2B right**). These ramp-based data, unlike pulse-based data, are consistent with the expected increase in sensitivity due to inflammation. Furthermore, injecting PF-24 and PF-71 significantly increased withdrawal latency in ramp-based tests (*T*_8_ = -5.646, *p* < 0.001, paired *t*-test).

**FIGURE 2.**
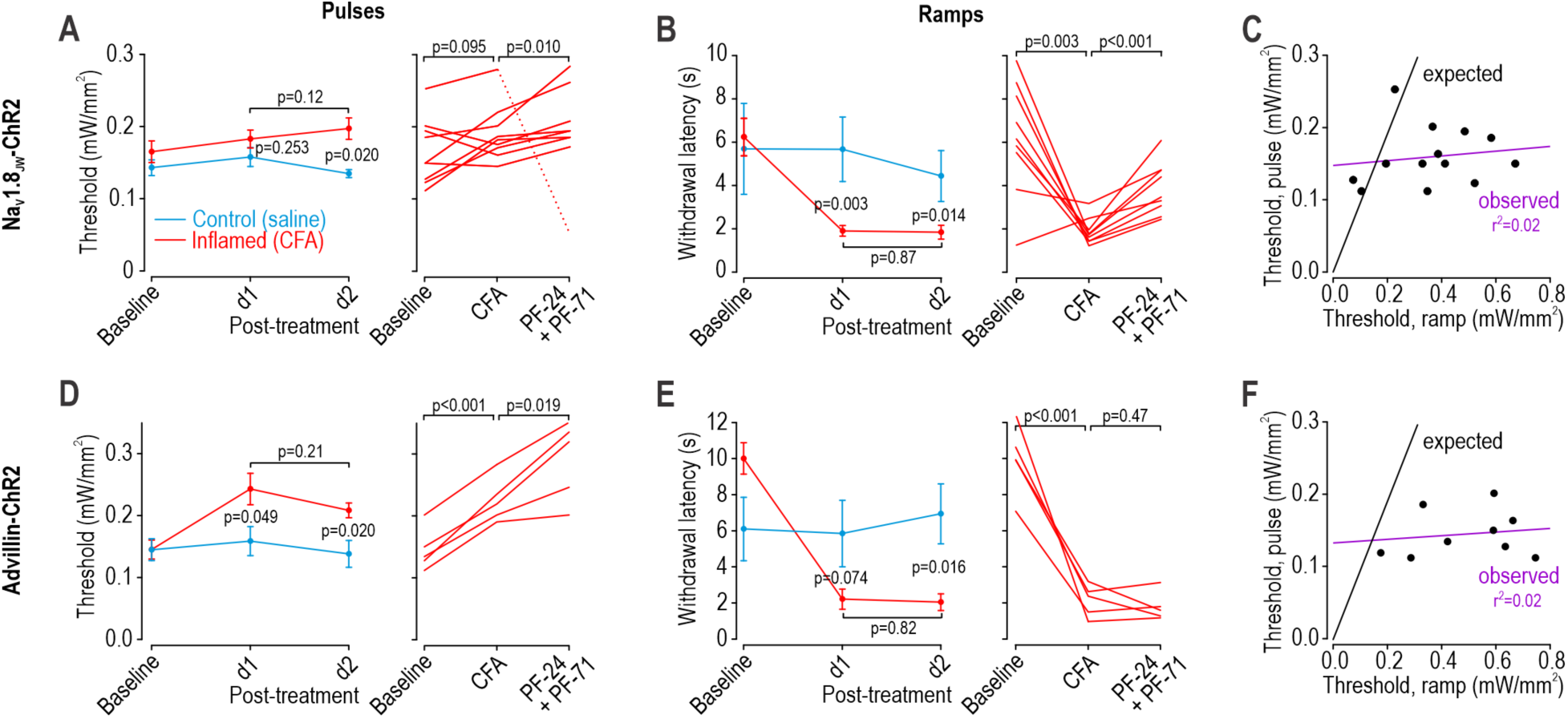
Comparison of pain sensitivity measured by pulsed and ramped photostimulation. **(A-C)** Testing in Na_V_1.8_JW_-ChR2 mice, which express ChR2 predominantly in nociceptors. *N* = 9 mice injected with CFA (red). *N* = 4 mice injected with saline (blue) (**A**) Pulse-based threshold measurements. Left: Threshold did not differ between groups at baseline (*T*_11_ = 0.922, *p* = 0.38, unpaired *t*-test) or one day post-injection (*T*_11_ = 1.206, *p* = 0.25), but was higher in the CFA group by day 2 (*T*_11_ = 2.712, *p* = 0.020). The difference between days 1 and 2 was not significantly different within the CFA group, (*T*_8_ = -1.733, p = 0.12, paired t-test) and data were therefore pooled. Right: The modest CFA-induced increase from baseline was not significant (*T*_8_ = -1.892, *p* = 0.095, paired t-test). Subsequent application of PF-24 and PF-71 caused a significant increase (*T*_7_ = -3.502, *p* = 0.010) after exclusion of one outlier (dashed line). (**B**) Ramp-based latency measurements. Left: Withdrawal latency did not differ between groups at baseline (*T*_11_ = 0.291, *p* = 0.78, unpaired *t*-test). Latency in the CFA group was significantly shorter than in controls on day 1 (*T*_11_ = -3.734, *p* = 0.003) and day 2 (*T*_11_ = 2.937, *p* = 0.014). The difference between days 1 and 2 was not significantly different within the CFA group, (*T*_8_ = -0.163, *p* = 0.88, paired *t*-test) and data were therefore pooled. Right: The CFA-induced reduction from baseline was significant (*T*_8_ = -4.280, *p* = 0.003, paired *t*-test). Subsequent application of PF-24 and PF-71 caused a significant increase (*T*_7_ = -5.646, p < 0.001). (**C**) Animal-by-animal correlation between pulse- and ramp-based measurements. For ramp-based data, latency was converted to threshold based on the ramp intensity at the time of withdrawal; this is a linear re-scaling, which means comparing pulse-based thresholds with ramp-based latencies or thresholds yields equivalent correlations. There was no correlation (purple line, *r*^2^ = 0.02); black line shows expected relationship for comparison. **(D-F)** Testing in advillin-ChR2 mice, which express ChR2 in all afferent types. *N* = 5 mice injected with CFA (red). *N* = 4 mice injected with saline (blue). (**D**) Pulse-based threshold measurement. Left: Threshold did not differ between groups at baseline (*T*_7_ = 0.0048, *p* = 1.00, unpaired *t*-test) but threshold was significantly higher in the CFA group one day (*T*_7_ = 2.377, *p* = 0.049) and two days (*T*_7_ = 3.009, *p* = 0.02) post-injection. The difference between days 1 and 2 was not significantly different within the CFA group, (*T*_4_ = 1.512, *p* = 0.21, paired *t*-test) and data were therefore pooled. Right: The CFA-induced increase from baseline was significant (*T*_4_ = -11.243, *p* < 0.001, paired t-test), as was the increase caused by PF-24 and PF-71 (*T*_4_ = -3.803, *p* = 0.019). (**E**) Ramp-based latency measurements. Left: The longer latency in CFA mice compared with control mice was noticeable, but not significant (*T*_7_ = 2.133, *p* = 0.070, unpaired *t*-test). This difference reversed one day post injection (*T*_7_ = -2.097, *p* = 0.074) and was significant by day 2 (*T*_7_ = -3.159, *p* = 0.016). The difference between days 1 and 2 was not significantly different within the CFA group, (*T*_4_ = 0.257, *p* = 0.81, paired *t*-test) and data were therefore pooled. Right: The CFA-induced reduction from baseline was significant (*T*_4_ = 7.790, *p* = 0.001, paired *t*-test) but subsequent application of PF-24 and PF-71 failed to cause a significant increase (*T*_4_ = 0.791, *p* = 0.047). (**F**) Animal-by-animal correlation between pulse- and ramp-based measurements. There was no correlation (*r*^2^ = 0.02).

To compare pulses and ramps, ramp-based latency data were converted to thresholds based on the light intensity (i.e. ramp level) at the time of withdrawal, and the resultant threshold was plotted against the pulse-based threshold measured in the same mouse. If afferents are equivalently activated by pulses and ramps, the correlation should be strong (notwithstanding within-mouse variability) and have a slope of one. If afferent activation differs quantitatively between pulses and ramps (e.g. if activation depends on the integral of ChR2 activation, and integration time differs between pulses and ramps), then the slope should deviate from one but correlation should remain high. However, if afferent activation differs qualitatively between pulses and ramps (e.g. because different afferent types are preferentially activated, or are activated with different synchrony), the correlation should break down. Using baseline data pooled across conditions, we found no correlation between pulse- and ramp-based thresholds (**Fig. 2C**), suggesting the two photostimulus waveforms activate afferents in distinct ways.

Parallel testing with pulses and ramps was also conducted on Advillin-ChR2 mice, which express ChR2 in all afferents. Contrary to expectations, but consistent with Na_V_1.8_JW_-ChR2 mice, threshold measured with pulses was higher in inflamed mice compared to controls (day 1: *T*_7_ = -2.377, *p* = 0.049; day 2: *T*_7_ = -3.009, *p* = 0.020; unpaired *t*-tests) and inflammation caused a significant increase in threshold relative to baseline (*T*_4_ = -11.243, *p* < 0.001, paired *t*-test) (**Fig. 2D**). On the other hand, latency measured with ramps was shorter in inflamed mice compared to controls (day 1: *T*_7_ = 2.097, *p* = 0.074; day 2: *T*_7_ = 4.896, *p* = 0.016, unpaired *t*-tests) and inflammation significantly reduced latency relative to baseline (*T*_4_ = 7.790, *p* = 0.001, paired *t*-test) (**Fig. 2E**). Subsequent injection of PF-24 and PF-71 did not significantly increase withdrawal latency (*T*_4_ = 0.791, *p* = 0.47, paired *t*-test) despite causing a significant increase in the pulse-based threshold (*T*_4_ = -3.803, *p* = 0.019). Like in Na_V_1.8_JW_-ChR2 mice, threshold values determined by pulses and ramps were not correlated (**Fig. 2F**).

To summarize, the expected CFA-induced decrease in threshold was not seen in pulse-based testing in either Na_V_1.8_JW_-ChR2 or Advillin-ChR2 mice; if anything, threshold was slightly increased. In contrast, the expected CFA-induced decrease in latency was observed with ramp-based testing. This discrepancy hints that different photostimulus waveforms activate primary afferents in qualitatively distinct ways (see Introduction), consistent with the lack of correlation (within the same animal) between pulse- and ramp-based threshold measurements. One should also consider the practicalities of measuring sensitivity with each waveform: We averaged three measurements per mouse per session for each testing protocol, but each determination of threshold required a sequence of 5.8 ± 0.9 (mean ± SD) pulses titrated with an up-down method, whereas each measurement of withdrawal latency involved a single ramp. Pulse-based testing therefore takes longer and entails many more stimuli, thus increasing the risk of learning, stress, etc.

No sex differences were observed. In Na_V_1.8_JW_-ChR2 mice, withdrawal latency at baseline in males (7.123 ± 0.963 s, mean ± SEM, *n* = 7) and females (4.826 ± 1.317, *n* = 6) was not significantly different (*T*_11_ = 1.436, *p* = 0.18, unpaired *t*-test), nor did the inflammation-induced change in latency differ between males (-4.486 ± 1.176 s, *n* = 5) and females (-4.213 ± 1.981 s, *n* = 4) (*T*_7_ = -0.125, p = 0.90, unpaired *t*-test). In Advillin-ChR2 mice (including additional mice not reported in Figure 2), latency at baseline in males (7.246 ± 0.563 s, *n* = 28) and females (6.146 ± 0.981, *n* = 12) was not significantly different (*T*_38_ = 1.027, *p* = 0.31, unpaired *t*-test); too few mice were tested with CFA to subdivide by sex. In Na_V_1.8_RK_-ChR2 mice reported below, latency at baseline in males (2.278 ± 0.250 s, *n* = 18) and females (2.310 ± 0.382, *n*= 8) was not significantly different (*T*_24_ = -0.0714, *p* = 0.94), nor did the inflammation-induced change in latency differ between males (-0.818 ± 0.256 s, *n* = 14) and females (-1.024 ± 0.408 s, *n* = 6) (*T*_18_ = 0.455, *p* = 0.65). Despite no sex differences being identified here, for behavioral tests conducted in the morning (shortly after lights on), sex differences have been observed with equivalent ramp-based testing at night [8].

### Altered ramp-evoked withdrawal in Na_V_1.8-ChR2 mice suggest hyperexcitable nociceptors

Using Na_V_1.8_RK_-ChR2 mice, we conducted additional ramp-based optogenetic testing. **Videos 1** and **3** show responses in the same mouse to ramped photostimulation before and 1 day after CFA. We found a significant interaction between treatment (saline vs CFA) and time (baseline vs. post-injection) (*F*_1,24_ = 10.914, *p* = 0.003, two-way repeated measures ANOVA). Withdrawal latency differed significantly between CFA and saline post-injection (*p* = 0.010, Student-Neuman-Keuls test) but not at baseline (*p* = 0.56). Withdrawal latency dropped significantly after CFA (**Fig. 3A left**, *p* < 0.001) whereas it increased slightly after saline (**Fig. 3A right**, *p* = 0.18). Based on within-mouse measurements, the 0.873 ± 0.212 s reduction after CFA differed significantly from the 0.519 ± 0.291 s increase after saline (**Fig. 3B**, *T*_24_ = -3.304, *p* = 0.003, unpaired *t*-test), revealing a large effect of inflammation (Hedges’ *g* = 1.54). These mice were either sacrificed for in vitro electrophysiology or were used to test effects of subtype-specific Na_V_ channel blockade.

**FIGURE 3.**
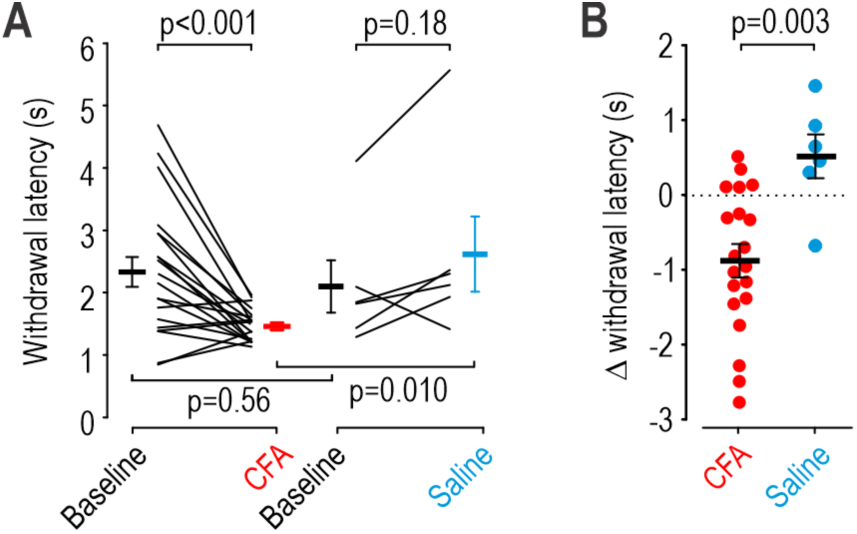
Effects of CFA on paw withdrawal from ramped photostimulation. (**A**) Withdrawal latency is plotted for each Na_V_1.8-ChR2 mouse at baseline and again after injecting CFA (left, *N* = 20 mice) or saline (right, *N* = 6 mice). Treatment (CFA vs saline) interacted significantly with time (baseline vs. post-injection) (*F*_1,24_ = 10.914, *p* = 0.003, two-way repeated measures ANOVA). Withdrawal latency (mean ± SEM) differed between CFA and saline conditions post-injection (1.472 ± 0.058 s vs 2.616 ± 0.606 s, *p* = 0.010, Student-Neuman-Keuls post-hoc test) but not at baseline (2.345 ± 0.239 s vs 2.097 ± 0.420 s, *p* = 0.56). (**B**) Change in latency due to treatment calculated within each mouse. The reduction in withdrawal latency due to CFA (red, -0.873 ± 0.212 s) differed significantly from the increase due to saline (blue, 0.519 ± 0.291 s) (*T*_24_ = -3.304, *p* = 0.003, unpaired *t*-test).

Inflammation reduced ramp-based withdrawal latencies in all three mouse lines tested, but those mouse lines responded differently at baseline. Those differences are considered later (see Fig. 7).

First, we sought to explore the neural correlates of the withdrawal behavior observed in Na_V_1.8_RK_-ChR2 mice, especially the changes due to inflammation. Because optogenetic stimulation bypasses natural transduction, inflammation-induced changes in mechanical or thermal transduction should not contribute to altered optogenetic sensitivity; instead, optogenetic testing helps isolate effects of altered excitability in ChR2-expressing afferents. That said, central sensitization could allow equivalent optogenetic activation of ChR2-expressing afferents to drive faster withdrawal. Thus, to rule in peripheral sensitization, our next step was to measure how inflammation affects the excitability of ChR2-expressing afferents, and how such changes impact responses to ramped photostimulation. We also ruled out effects of inflammation on ChR2 expression or gating so that changes in optogenetic responses can be confidently ascribed to altered excitability.

### Inflammation caused increased somatic excitability consistent with upregulation of Na_V_1.7

To explore inflammation-induced changes in nociceptor excitability, we electrophysiologically tested small-diameter ChR2-positive neurons harvested from control and inflamed (CFA-injected) Na_V_1.8_RK_-ChR2 mice. All recordings were conducted 2-8 hours after dissociation to avoid changes induced by culturing [54; 56]. Neurons from CFA-injected mice (henceforth labeled “inflamed” neurons) were excited by weaker current injection and fired more vigorously than control neurons (**Fig. 4A**). Rheobase (i.e. the weakest current required to evoke a spike) in control neurons (14.5, 6 – 28 pA; median, interquartile range [IQR]) was higher than in inflamed neurons (4, 2 – 8.75 pA) (*U*_27_ = 36, *p* = 0.003, Mann-Whitney test) (**Fig. 4B**). Evoked firing rates revealed a significant interaction between stimulus intensity and condition (*F*_6,168_ = 4.834, *p* < 0.001, two-way repeated measures ANOVA) (**Fig. 4C**). Because rheobase was lower after inflammation and because we incremented stimulus intensity as multiples of rheobase, weaker stimuli were applied to inflamed neurons than to control neurons, which means even stronger effects of inflammation would have been seen by matching absolute stimulus intensities across conditions. The difference in evoked firing rate was at least partly due to the higher proportion of transient spiking neurons (i.e. unable to ever fire >2 spikes) relative to repetitive spiking neurons in control vs. inflamed conditions (χ^2^ = 4.55, *p* = 0.033) (**Fig. 4C inset**). Several other metrics were unchanged between control and inflamed neurons: Resting membrane potential (-64.1 ± 2.5 mV vs -61.6 ± 1.8 mV, *T*_27_ = -0.833, *p* = 0.412), input resistance (2.29 ± 0.27 MΩ vs 2.35 ± 0.22 MΩ, *T*_27_ = -0.164, *p* = 0.871), and capacitance (15.45 ± 1.05 pF vs 14.90 ± 0.93 pF, *T*_27_ = 0.399, *p* = 0.693). These results show that inflammation increased the somatic excitability of nociceptors without affecting passive membrane properties.

**FIGURE 4.**
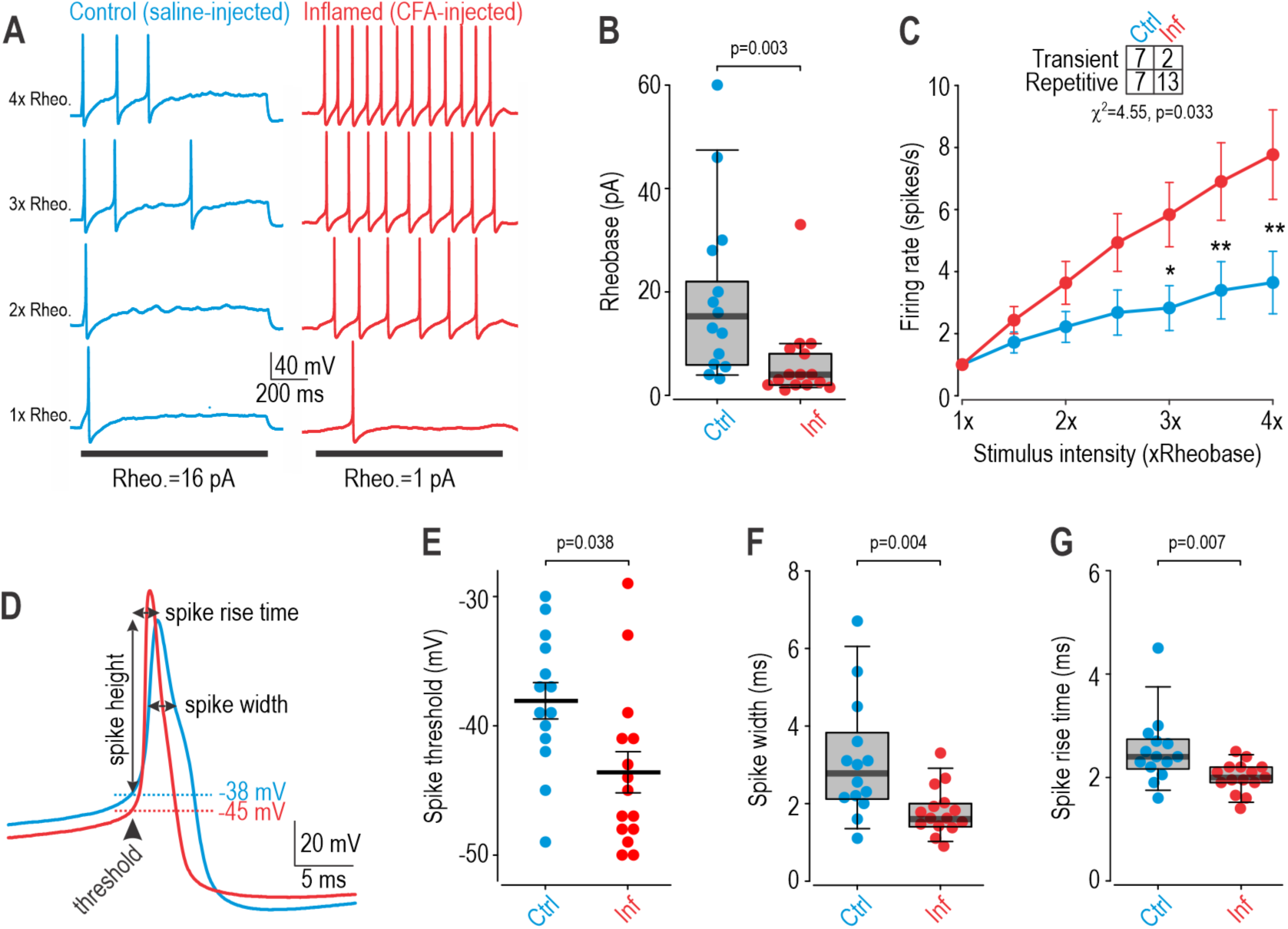
Effects of CFA on the somatic excitability of nociceptors. *N* = 14 control neurons (blue) and 15 inflamed neurons (red) for all panels. (**A**) Sample current clamp recordings show spiking evoked by current injection in a typical control (blue) and inflamed (red) neuron. Rheobase was determined for each neuron as the weakest current step required to evoke a spike. Current injection was incremented as multiples of rheobase for each neuron. Inflamed neurons responded vigorously even to weak stimulation. (**B**) Rheobase was significantly lower in inflamed neurons (4, 2 – 8.75 pA; median, IQR) than in control neurons (14.5, 6 – 28 pA) (*U*_27_ = 36, *p* = 0.003, Mann-Whitney *U*-test). (**C**) Evoked firing rates depended jointly on stimulus intensity and condition (interaction, *F*_6,168_ = 4.834, *p* < 0.001, two-way repeated measures ANOVA). Results of post-hoc Student-Neuman-Keuls tests are summarized on graph (*, *p* < 0.05; **, *p* < 0.01). Weaker stimuli were applied to inflamed neurons than to control neurons because of the difference in rheobase, meaning an even a larger inflammation-induced increase in evoked firing would be seen if absolute stimulus intensities was matched. Table shows that transient spiking neurons (i.e. neurons that never fire >2 spikes during current injection) are significantly less common after inflammation than in control conditions (χ^2^ = 4.55, *p* = 0.033). (**D**) Enlarged traces highlight differences in spike waveform. Arrows show measurements compared in panels E-G. The first spike evoked by rheobasic stimulation was measured. (**E**) Spike threshold was significantly more negative in inflamed neurons (-43.6 ± 1.6 mV) than in control neurons (-38.1 ± 1.4 mV) (*T*_27_ = 2.58, *p* = 0.016, unpaired *t*-test). Threshold was defined as the voltage where dV/dt exceeds 5 mV/ms (**F**) Spikes were significantly narrower in inflamed neurons (1.6, 1.413 – 1.975 ms) than in control neurons (2.775, 2.15 – 3.6 ms) (*U*_27_ = 39, *p* = 0.004). Width was measured at half of the spike height. (**G**) Spikes rose significantly faster in inflamed neurons (2.0, 1.9 – 2.2 ms) than in control neurons (2.4, 2.2 – 2.7 ms) (*U*_27_ = 42.5, *p* = 0.007). Rise time was measured from threshold crossing to peak voltage.

Additional changes in active membrane properties were revealed by comparing the spike waveform in control and inflamed neurons (**Fig. 4D**). Spike threshold in control neurons (-38.1 ± 1.4 mV) was significantly more depolarized (i.e. less negative) than in inflamed neurons (-43.6 ± 1.6 mV) (*T*_27_ = 2.58, *p* = 0.016, unpaired t-test) (**Fig. 4E**) which translated to a significant difference in spike height (86.25, 84 – 89 mV vs 91, 90.125 – 96.25 mV; *U*_27_ = 57, *p* = 0.038). Conversely, spike width in control neurons (2.775, 2.15 – 3.6 ms) was greater than in inflamed neurons (1.6, 1.413 – 1.975 ms) (*U*_27_ = 39, *p* = 0.004) (**Fig. 4F**). Spike rise time in control neurons (2.4, 2.2 – 2.7 ms) was also greater than in inflamed neurons (2.0, 1.9 – 2.2 ms) (*U*_27_ = 42.5, *p* = 0.007) (**Fig. 4G**). These changes, consistent with previous data [19], reflect a shift in the dominant sodium channel isoform from Na_V_1.8 to Na_V_1.7 after inflammation [54], which has implications for drug efficacy (see Fig. 6).

**FIGURE 5.**
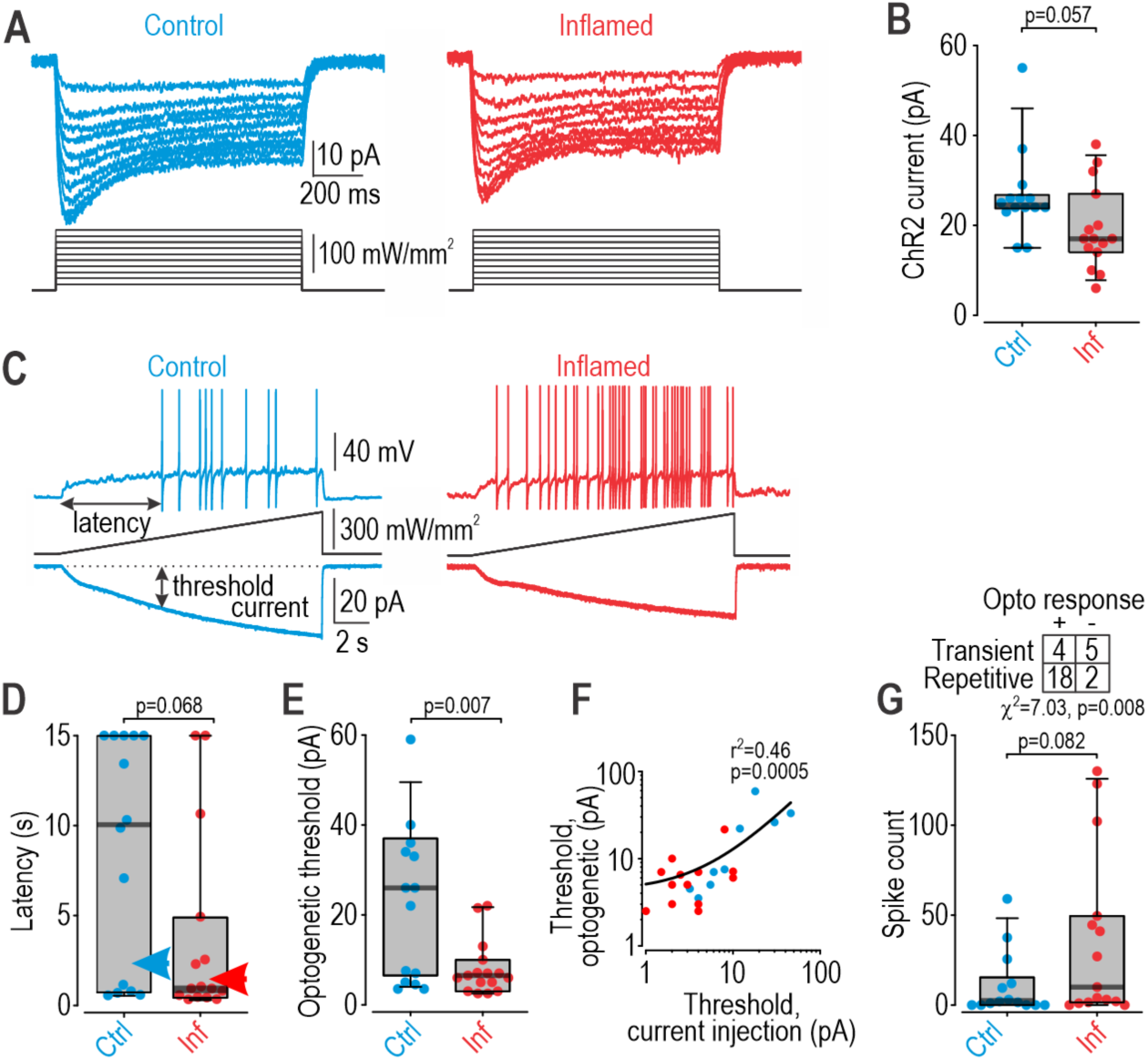
Effects of CFA on ChR2-expression and optogenetically evoked somatic spiking. The same neurons tested electrophysiologically in Figure 4 were also stimulated optogenetically. **(A)** Sample voltage clamp recordings show current evoked by optogenetic stimulation with blue light applied as 1 s-long step at different intensities to a typical control (blue) and inflamed (red) neuron. No differences in ChR2 gating kinetics were observed. **(B)** ChR2 current evoked by 156 mW/mm^2^ blue light in inflamed neurons (17, 14.25 – 25.25 pA) was smaller than in control neurons (24.5, 24 – 26 pA) but the difference did not reach significance (*U*_27_ = 61, *p* = 0.057). **(C)** Sample current clamp recordings (top) show spiking evoked by a 15-s-long optogenetic ramp. Bottom traces show ChR2 current evoked by the equivalent optogenetic ramp applied during voltage clamp. Arrows highlight measurements compared in panels D-F. Sample traces in C are from the same neurons illustrated in panel A. Spiking was increased in inflamed neuron because of increased excitability, despite reduced ChR2 current. **(D)** Latency to first spike was significantly shorter in inflamed neurons (0.940, 0.489 – 4.330 s) than in control neurons (10.085, 0.785 – 15 s) (*U*_27_ = 64, *p* = 0.068). Neurons that did not respond with any spikes were treated as having a latency of 15 s. Arrowheads mark average withdrawal latency for respective condition. **(E)** Optogenetic threshold (ChR2 current at the time of the first spike) was significantly lower in inflamed neurons (6.5, 3.5 – 9.275 pA) than in control neurons (26, 7 – 36 pA) (*U*_27_ = 42.5, *p* = 0.007). **(F)** Optogenetic threshold and rheobase (threshold based on current injection) measured within the same cell were significantly correlated (r^2^ = 0.46, *p* = 0.0005). Data are plotted on log scales to distribute data points more evenly for visualization. **(G)** The number of optogenetically evoked spikes in inflamed neurons (10, 1.625 – 48.25 spikes) was higher than in control neurons (1.75, 0 – 12 spikes), but the difference did not reach significance (*U*_27_ = 65, *p* = 0.082). Table shows that repetitive spiking neurons are significantly more likely than transient spiking neurons to respond to the optogenetic ramp with at least one spike (χ^2^ = 7.03, *p* = 0.008), which is notable given the shift from transient to repetitive spiking after inflammation (see table on Fig.3C).

**FIGURE 6.**
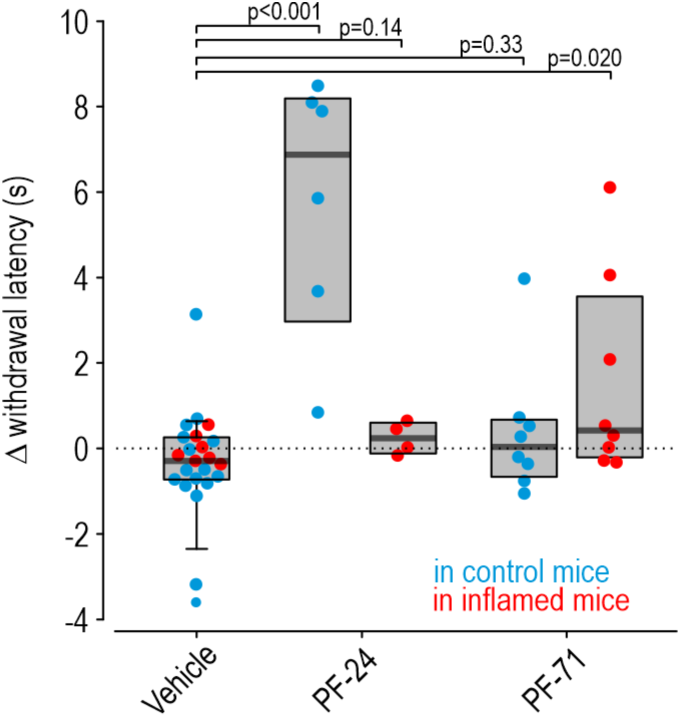
Differential effects of subunit-selective Na_V_ inhibitors on withdrawal confirm that axons experience an inflammation-induced shift from Na_V_1.8 to Na_V_1.7. Data are plotted as the change in withdrawal latency induced by injecting drug or vehicle into the optogenetically stimulated paw. Injecting vehicle had an equivalent effect in control and inflamed mice (*U*_21_ = 34, *p* = 0.15, Mann-Whitney *U* test) and data were therefore pooled to give a median latency change of -0.290, -0.722 – 0.239 s. The effect of blocking Na_V_1.8 or Na_V_1.7 with PF-24 and PF-71, respectively, are compared against the vehicle effect. PF-24 significantly increased withdrawal latency by 6.871, 3.676 – 8.096 s in control mice (*U*_27_ = 1, *p* < 0.001) but caused only a 0.239, -0.073 – 0.552 s increase in inflamed mice (*U*_25_ = 24, *p* = 0.14). Conversely, PF-71 increased withdrawal latency by only 0.039, -0.564 – 0.624 s in control mice (*U*_29_ = 70, *p* = 0.33) but significantly increased it by 0.417, -0.133 – 3.066 s in inflamed mice (*U*_29_ = 40, *p* = 0.020).

**FIGURE 7.**
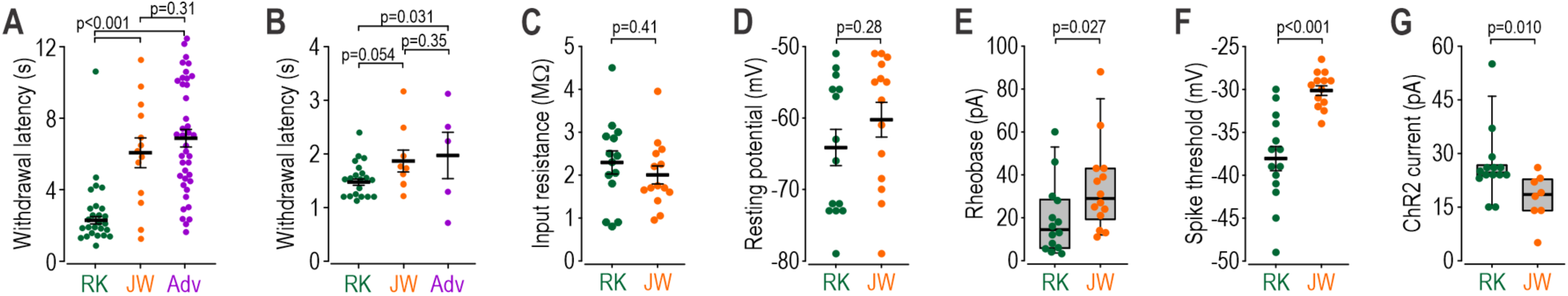
Comparison across mouse lines. (**A**) Ramp-based withdrawal latencies at baseline. Baseline data from all mice regardless of subsequent testing were included. *N* = 26 NaV1.8_JK_-ChR2 mice (RK, green). *N* = 13 NaV1.8_JW_-ChR2 mice (JW, orange). *N* = 40 Advillin-ChR2 mice (Adv, purple). Genotype had a significant effect (F_2,76_ = 25.823, one-way ANOVA). Results of post-hoc Student-Newman-Keuls tests are indicated on the graph. (**B**) Ramp-based withdrawal latencies after CFA (using average across day 1 and 2 post-injection). *N* = 20, 9, and 5 mice from left to right. Genotype had a diminished but still significant effect (F_2,31_ = 4.535, one-way ANOVA). Results of post-hoc Student-Newman-Keuls tests are indicated on the graph. **(C-G)** Electrophysiological data from control JW mice (N = 14 small-diameter neurons unless otherwise indicated) for comparison with data from RK mice (N=14 cells, from Figures 2 and 3). Data are summarized as mean ± SEM or with a boxplot (median and interquartile range) depending on the distribution; an unpaired *t*-test or Mann-Whitney test was applied as appropriate. (**C**) Input resistance did not differ between RK and JW neurons (*T*_26_ = 0.845, *p* = 0.41, unpaired *t*-test). (**D**) Resting membrane potentials did not differ between RK and JW neurons (*T*_26_ = -1.108, *p* = 0.28, unpaired *t* -test). (**E**) Rheobase was significantly higher in JW neurons (*U*_26_ = 49.5, *p* = 0.027, Mann-Whitney test). (**F**) Spike threshold was also significantly less negative (more depolarized) in JW neurons (*T*_26_ = -5.234, *p* < 0.001, unpaired *t* -test). (**G**) ChR2 current evoked by 156 mW/mm^2^ of blue light was significantly smaller in JW neurons (*U*_20_ = 18, *p* = 0.010, Mann-Whitney test; *N* = 8 JW neurons).

### Hyperexcitability increases optogenetically evoked spiking consistent with behavioral changes

Next, we checked if inflammation affects ChR2 function or expression. Voltage clamp recordings of the current evoked by different photostimulus intensities applied as 1 s-long steps exhibited no differences in ChR2 gating between control and inflamed neurons but suggested a modest reduction in ChR2 expression (**Fig. 5A**). The ChR2 current evoked by 156 mW/mm^2^ blue light in control neurons (24.5, 24 – 26 pA) was slightly larger than in inflamed neurons (17, 14.25 – 25.25 pA) but the difference did not reach significance (**Fig. 5B**, *U*_27_ = 61, *p* = 0.057). The modest reduction in ChR2 current did not offset the large increase in excitability in inflamed neurons according to optogenetically evoked spiking (see below).

**Figure 5C** illustrates sample responses to ramped photostimulation. Kinetics match the ramps used for behavioral testing but the intensity of light reaching the target cannot be directly compared between in vitro and in vivo conditions, and nor can somatic and axonal excitability be directly compared. Nevertheless, latency to the first optogenetically evoked spike in control neurons (10.085, 0.785 – 15 s) was much longer than in inflamed neurons (0.940, 0.489 – 4.330 s) (**Fig. 5D**, *U*_27_ = 63, *p* = 0.068). A subset of neurons did not respond with any spikes and were treated as having a latency of 15 s (i.e. the duration of the ramp). By measuring in voltage clamp the ChR2 current evoked in the same neuron to an equivalent photostimulus ramp, threshold ChR2 current was identified as the ChR2 current at the time of the first spike (see arrows on Fig. 5C). Threshold ChR2 current was significantly higher in control neurons (26, 7 – 36 pA) than in inflamed neurons (6.5, 3.5 – 9.275 pA) (**Fig. 5E**, *U*_27_ = 42.5, *p* = 0.007). Plotting each neuron’s rheobase (based on current injection) against its optogenetic threshold (based on ChR2 current) revealed a significant correlation (**Fig. 5F**, r^2^ = 0.46, *p* = 0.0005). Fewer spikes were evoked in control neurons (1.75, 0 – 12 spikes) than in inflamed neurons (10, 1.625 – 48.25 spikes), but this difference did not reach significance (**Fig. 5G**, *U*_27_ = 65, *p* = 0.082). Interestingly, neurons identified as transient spiking (based on their response to current injection, see above) were significantly less likely to respond with any spikes to ramped photostimulation (**Fig. 5G inset**, χ^2^ = 7.03, *p* = 0.008). This is consistent with transient spiking neurons requiring abrupt depolarization to initiate spikes [37; 38]; photostimulus ramps (unlike pulses) evidently do not cause rapid enough depolarization to evoke spikes in most transient spiking neurons, which is notable since a higher proportion of neurons spike repetitively after inflammation (see inset on Fig. 4C),

Notwithstanding limitations in quantitatively comparing optogenetically evoked spiking with optogenetically evoked paw withdrawal (see above), consider that 60% of inflamed cells (9 of 15) were activated before withdrawal occurred in inflamed Na_V_1.8_RK_-ChR2 mice (1.472 s) whereas only 36% of control cells (5 of 14) were activated before withdrawal occurred in saline-injected mice (2.616 s). This suggests that inflamed cells, because of their heightened excitability, were recruited by weaker optogenetic stimulation, thus firing and triggering withdrawal at shorter latency. By comparison, despite a near-significant difference in the total number of spikes evoked by the optogenetic ramp in each condition, if the count is restricted to spikes occurring *before* the average withdrawal latency in each condition, few spikes occur in control or inflamed neurons (0, 0 – 2 spikes vs 2, 0 – 4) (*U*_27_ = 72.5, *p* = 0.14). From this, it seems most likely that the number of activated neurons dictate when withdrawal is initiated, since most activated neurons each contribute only one spike by the time withdrawal occurs. Responses to pulsed photostimuli provide additional context [16]: Withdrawal from strong pulses occurred in as little as 20 ms but withdrawal to threshold pulses took as long as 500 ms, consistent with the ∼400 ms latency measured for pulses reported in Figure 2. Since optogenetic ramps approach the threshold amplitude slowly, one should consider stimulus amplitude 400 ms before the observed response (based on the 400 ms latency for just-threshold pulses). For the ramped photostimuli reported here, approximately 30% of neurons were activated 400 ms before the withdrawal latency in each condition, suggesting enough control neurons were recruited ∼2 s after the start of the optogenetic ramp (2.616 – 0.4 = 2.216 s) whereas inflamed neurons required half as long to be sufficiently recruited (1.472 – 0.4 = 1.072 s), implying they required half as much stimulation because of their heightened excitability. These calculations are speculative but provide a framework to guide future experiments.

### Nociceptor axons experience the same shift in Na_V_ subunit contribution as somata

We showed previously that nociceptors rely on Na_V_1.8 channels under normal conditions but that Na_V_1.7 channels become more important after inflammation [54], consistent with spike waveform changes described in Figure 4D-G. Extrapolating from those data, we predicted that blocking Na_V_1.8 (with PF-24) would delay paw withdrawal in control mice but not in inflamed mice, whereas blocking Na_V_1.7 (with PF-71) would delay withdrawal in inflamed mice but not in control mice. In other words, the analgesic efficacy of PF-24 should decrease after inflammation whereas the efficacy of PF-71 should increase. Either drug was injected into the paw at the stimulation site to modulate axon excitability while measuring effects on withdrawal from ramped photostimulation.

**Figure 6** reports the change in withdrawal latency produced by injecting drug or vehicle (saline-diluted DMSO) into control or inflamed mice. Vehicle produced a modest reduction in withdrawal latency in control mice (0.583, -0.845 – 0.217 s) and inflamed mice (0.147, -0.275 – 0.225 s); the effect did not differ between groups (*U*_21_ = 34, *p* = 0.15, Mann-Whitney test) and data were therefore pooled to give a median latency change of -0.290, -0.722 – 0.239 s against which to compare drug effects. As predicted, PF-24 significantly increased withdrawal latency by 6.871, 3.676 – 8.096 s in control mice (*U*_27_ = 1, *p* < 0.001; Hedges’ *g* = 3.53) while insignificantly increasing withdrawal latency by 0.239, -0.073 – 0.552 s in inflamed mice (*U*_25_ = 24, *p* = 0.14; Hedges’ *g* = 0.49). Conversely, PF-71 insignificantly increased withdrawal latency by 0.039, -0.564 – 0.624 s in control mice (*U*_29_ = 70, *p* = 0.33; Hedges’ *g* = 0.54) but significantly increased it by 0.417, -0.133 – 3.066 s in inflamed mice (*U*_29_ = 40, *p* = 0.020; Hedges’ *g* = 1.19). Effect sizes (quantified as Hedges’ *g* values) showed that blocking either Na_V_ isoform delayed withdrawal in each condition, but blocking Na_V_1.8 had ∼7× greater effect in control mice (3.53/0.49 = 7.20) whereas blocking Na_V_1.7 had ∼2× greater effect in inflamed mice (1.19/0.54 = 2.20). These results confirm that nociceptor axon terminals experience a similar inflammation-induced shift in Na_V_ expression as somata.

### Neural correlates of differences in optogenetic sensitivity across mouse lines

The difference in withdrawal latencies between mouse lines (see Figures 2 and 3) merited consideration. Comparing withdrawal latency at baseline in Na_V_1.8_RK_-ChR2 mice (2.288 ± 0.205 s), Na_V_1.8_JW_-ChR2 mice (6.063 ± 0.832 s), and Advillin-ChR2 mice (6.916 ± 0.491 s) revealed a significant effect of genotype (*F*_2,76_ = 25.823, *p* < 0,001, one-way ANOVA) with Na_V_1.8_RK_-ChR2 mice differing significantly from the other two lines (*p* < 0.001, Student-Newman-Keuls tests) whereas Na_V_1.8_JW_-ChR2 and Advillin-ChR2 mice were similar (*p* = 0.31) (**Fig. 7A**). After inflammation, a small but still significant effect remained (F_2,31_ = 5.535, *p* = 0.019), with the latency in Na_V_1.8_RK_-ChR2 mice (1.472 ± 0.058 s) still significantly shorter than in Advillin-ChR2 mice (2.127 ± 0.398 s, *p* = 0.031) and nearly so in Na_V_1.8_JW_-ChR2 mice (1.866 ± 0.203 s, *p* = 0.054) (**Fig. 7B**).

To explore the basis for baseline differences between Na_V_1.8_RK_-ChR2 and Na_V_1.8_JW_-ChR2 mice (henceforth labeled RK and JW mice), we conducted additional electrophysiological testing of ChR2-expressing small-diameter neurons from control JW mice for comparison with data already reported for control RK mice (see Fig. 4). Input resistance (RK: 2.292 ± 0.271 MΩ; JW: 2.004 ± 0.209 MΩ; n=14 cells per group) was not significantly different (*T*_26_ = 0.845, *p* = 0.41, unpaired t-test) (**Fig. 7C**) nor was resting membrane potential (RK: -64.1 ± 2.5 mV; JW: -60.3 ± 2.4 mV; *T*_26_ = -1.108, *p* = 0.28) (**Fig. 7D**). On the other hand, rheobase was significantly lower in RK mice (RK: 14.5, 6 – 28 pA; JW: 29, 21-43 pA; *U*_26_ = 49.5, *p* = 0.027, Mann-Whitney test) (**Fig. 7E**), concordant with their more negative spike threshold (RK -38.1 ± 1.4 mV; JW, -30.1 ± 0.6 mV; *T*_26_ = -5.234, *p* < 0.001) (**Fig. 7F**). Despite this quantitative difference in excitability, the proportion of transient spiking neurons did not differ between RK and JW mice (RK: 50%, 7/14; JW: 36%, 5/14; χ^2^ = 0.583, *p* = 0.45). Additionally, ChR2 current was higher in RK mice (RK: 24.5, 24-26 pA; JW: 18.5, 14 - 22.5 pA; *U*_22_ = 18, *p* = 0.010) (**Fig. 7G**). The combination of weaker ChR2 expression and lower nociceptor excitability in JW mice likely accounts, at least in part, for their slower withdrawal from ramped photostimulation compared with RK mice. Indeed, ramped photostimulation evoked spikes in only 3 of 14 neurons from control JW mice, which is a significantly lower proportion than the 9 out of 14 neurons activated from control RK mice (χ^2^ = 5.25, *p* = 0.022). Furthermore, the average first spike latency in JW neurons was 6.7 s, which aligns well with the 6.063 s withdrawal latency at baseline in those mice. We did not test inflamed neurons from JW mice. Testing various conditions in a single mouse line avoids effects of genotype, but such technical details should be kept in mind.

## DISCUSSION

We applied transcutaneous optogenetic stimulation to behaving mice to explore the potential of this stimulus modality to quantify changes in pain sensitivity, especially to help isolate the functional consequences of changes occurring in specific types of afferents. Although optogenetic stimulation is unnatural, it enables selective activation of afferents based on their expression of actuators like ChR2. Moreover, light has numerous benefits as a stimulus insofar as it can be applied from a distance with precise spatial and temporal control. Using new technology designed reproducibly stimulate awake behaving mice while precisely measuring their withdrawal (**Fig. 1**), we tested photostimulus pulses and ramps, and measured the threshold and latency to each photostimulus waveform (**Fig. 2**). Back-to-back comparison of the testing protocols favors ramp-based testing but also suggests that afferent activation differs between photostimulus waveforms. Electrophysiological testing in vitro confirmed that nociceptor excitability is increased by inflammation (**Fig. 4**) and that this hyperexcitability (rather than changes in ChR2) likely accounts for inflamed mice withdrawing more rapidly than naïve mice from ramped photostimuli (**Fig. 5**). Comparing withdrawal latencies with ramped-evoked spiking further suggests that withdrawal latency depends on the number of activated neurons, as opposed to changes in the firing rate of a fixed number of neurons. Results also suggest that inflammation shifts the Na_V_1.8:Na_V_1.7 ratio in favor of Na_V_1.7 (**Fig. 6**), consistent with changes measured in the soma.

Pain behavior testing is notoriously irreproducible [13] but improved methods can change this. Optogenetics allows for exquisite control of *which* neurons are activated, but steps must nevertheless be taken to control the strength and pattern of their activation (see Introduction). And so too must the behavioral consequences be accurately measured. Together, these technical improvements should reduce response variability, but they will not remove variability altogether; the stimulus-response relationship is still impacted by variations in stress, circadian rhythms, etc. (i.e. biological “noise”), which is why acclimation, time of testing, and countless other factors are so important [42]. Reducing noise – in the stimulus, measurement, or biological system – allows one to resolve smaller effects, or to resolve the same effect using fewer animals. Resolution also depends on the testing protocol: When varying stimulus intensity to measure threshold, a smaller step size yields more precise measurement and, in turn, finer resolution of differences between conditions, but it comes at the expense of needing to test more steps. Difficulty measuring responses often translates to responses being scored as present or absent, but the binary distinction is very subjective, and subtle (but potentially important) variations in responses are invariably glossed over. These limitations compound.

Manual application of photostimuli leads to variations in the intensity of light reaching the target. This causes photostimulus intensity to vary from one pulse to the next, or to vary over the course of a prolonged stimulus [16]. Robotic stimulation enables prolonged photostimulation at a consistent intensity, which makes it feasible to apply photostimulus ramps reproducibly and makes it worth measuring the associated withdrawal latency. Ramp-based testing of withdrawal latency resembles the Hargreaves test whereas pulse-based testing of threshold is more akin to von Frey testing with different filaments (although each filament should be applied gradually, which is unlike pulsed photostimuli). Using an efficient up-down strategy, each threshold measurement required a sequence of ∼6 pulses, whereas a latency measurement was acquired from each ramp. In that regard, ramp-based testing is far more efficient.

But the photostimulus waveform itself is likely to have other consequences (see Introduction). Ramp- and pulse-based threshold measurements were completely uncorrelated (see Figs. 2C and F), which suggests that each waveform (*i*) preferentially activates different ChR2-expressing neurons and/or (*ii*) activates neurons with different patterns. With respect to *i*, we found that photostimulus ramps tend not to activate neurons identified as transient spiking based on their response to current steps (see inset on Fig. 5G). With respect to *ii*, synchronized afferent input is important for somatosensation [43], and ramping photostimulus intensity avoids the grossly unnatural spike synchrony associated with pulses. One should recall that electrical pulses were once commonly used in pain research, but were eventually shown to be a poor way to measure sensitivity [27]. Responses are scored as present or absent for pulse-based testing whereas ramped-based testing requires accurate measurement of withdrawal latency, but such measurements are straightforward with our reflectance-based method (see Fig. 1B) and can also be achieved with high-speed videography [10]. Thus, strategically designed photostimulus waveforms applied reproducibly with the appropriate technology combined with accurate response measurements provide new opportunities for reproducible, quantitative optogenetic pain testing.

Many molecular changes are induced by inflammation [21; 25; 51]. The number of co-occurring changes and the variety of neuron types affected make it difficult to pinpoint which changes in which neurons contribute to altered pain sensitivity. By using Na_V_1.8-ChR2 mice, ChR2 should be preferentially expressed in nociceptors [3], thus enabling one to isolate the role of nociceptors in mediating withdrawal, and to measure the consequence of inflammatory changes in nociceptors. That said, Na_V_1.8-ChR2 mice also express ChR2 in C-low threshold mechanoreceptors; C-LTMRs are absent from glabrous skin but sparsely innervate a small patch of hairy skin between the pads [34] and could therefore be optogenetically activated. Some myelinated afferents also express ChR2 [46] and we cannot exclude their contribution. Only ∼10% of myelinated fibers express Na_V_1.8 in adulthood [4], so the expression pattern of ChR2 presumably reflects broader expression of Na_V_1.8 (and Cre recombinase) during development. That said, pharmacological blockade of Na_V_1.8 by PF-24 will predominantly affect C fibers, as will blockade of Na_V_1.7 by PF-71. Since optogenetic activation bypasses thermal and mechanical transduction, increased sensitivity to optogenetic stimulation after inflammation points to increased excitability, which we verified electrophysiologically (see Figs. 4 and 5). Changes in ChR2 expression and gating were excluded (see Fig. 5), but we cannot exclude that central sensitization did not also contribute to increased optogenetic sensitivity. Notably, whereas patch clamp recordings demonstrate changes in somatic excitability, our behavioral data point to inflammation-induced changes in the excitability of cutaneous terminals of ChR2-expressing afferents. Intradermal injection of PF-24 and PF-71 at the photostimulation site help further target changes in *axon* excitability.

Our observation that inflammation-induced changes in cutaneous axon excitability mediate an increase in pain sensitivity is supported by the effects of subtype-selective Na_V_ inhibitors injected into the paw. Extrapolating from past experiments on somatic excitability [54], we predicted that pain sensitivity in control mice depends mostly on Na_V_1.8, whereas pain sensitivity in inflamed mice depends mostly on Na_V_1.7. Such knowledge is obviously important for choosing the best drug for a given pain condition. Inferring which ion channel to target based on an observed excitability change is, however, complicated by the fact similar excitability changes can arise from distinct molecular changes [39–41]. Matters are further complicated by certain changes being unique to axons [49; 52], which risks them being overlooked in somatic recordings, but so too might somatic changes not generalize to axons. In the case of Na_V_ isoforms, both compartments evidently experience a similar inflammation-induced shift from Na_V_1.8 to Na_V_1.7. Beyond testing if a drug target on nociceptors changes, optogenetic testing can also help determine via which neuron type a drug modulates behavior, if, for example, the target is present on >1 type of neuron, especially if those neuron types are co-activated by natural stimuli (notwithstanding imperfect ChR2 expression; see above). Many other experiments are conceivable.

Overall, these results demonstrate the utility of optogenetic pain testing. The opportunities are enhanced by technology enabling reproducible stimulation and accurate response measurements, and by using photostimulus waveforms designed to optimize the sensitivity and efficiency of testing.

## Acknowledgements

We thank Jason Jeong for technical assistance and Stéphanie Ratté for feedback on the manuscript. This work was supported by a Foundation Grant from the Canadian Institutes of Health Research (FDN167276).

## Author contributions

The project was conceived by CD and SAP. Experiments were carried out by YFX with assistance from CD. Data were analyzed by YFX with assistance from CD and SAP. SAP supervised all research. SAP produced the final manuscript with input from YFX and CD.

